# The combinatorial effect of terminators and introns on the levels and stability of stable transgene expression in plants

**DOI:** 10.64898/2026.08.17.745381

**Authors:** Buddhini Ranawaka, Kylie Shand, Peter M. Waterhouse, Felipe F. de Felippes

## Abstract

Most transgene applications require high and sustained expression, particularly in stably transformed plants. Achieving optimal transgene performance, however, depends on the combined influence of multiple genetic and regulatory factors. In previous work, we systematically evaluated the contribution of different genetic elements to transient transgene expression and demonstrated that terminators are key determinants of transgene performance by reducing transcriptional read-through and preventing transgene silencing. Here, we extend these findings by investigating the roles of terminators and introns in the expression of transgenes in stably transformed plants. Our results show that optimal transgene performance arises from the complementary actions of these two elements. Terminator choice was a major determinant of transgene expression levels, whereas introns played a critical role in maintaining expression stability. We further demonstrate a strong relationship between transgene expression levels and small RNA accumulation and show that intron-containing endogenous genes are enriched among highly expressed and stress-responsive genes, suggesting that intron-mediated protection from silencing may facilitate higher levels of gene expression and have contributed to the emergence and evolutionary retention of intron-containing genes.

## Introduction

Transgenes are crucial for various applications in modern plant biology, including functional genomic studies, the utilisation of plants as biofactories, and the development of crop varieties with novel agronomic traits. The level of transgene expression is a key factor when working with transgenic systems. Most applications require that transgenes be highly expressed; accordingly, several molecular approaches exist to achieve high transgene expression. The most common strategies rely on increasing transcription and translation rates, usually by using strong promoters and translation enhancers, respectively (Streatfield, 2007; Feng *et al*., 2022). Nonetheless, there are other genetic elements, many of which are overlooked, that can significantly affect how transgenes are expressed. One of these elements is the gene terminator, a region of DNA located downstream of the coding sequence, whose primary function is associated with transcriptional termination and polyadenylation of the messenger RNA (mRNA) (Bernardes and Menossi, 2020; de Felippes and Waterhouse, 2022). In addition to its fundamental role in mRNA maturation, an ever-growing number of studies show that the choice of terminator can substantially affect the final levels of transgene expression, indicating a more direct role of this genetic element in controlling gene expression (Ingelbrecht *et al*., 1989; Marshall *et al*., 1997; Chen *et al*., 1998; Richter *et al*., 2000; Ali and Taylor, 2001a; Schünmann *et al*., 2003; Nagaya *et al*., 2009; Yang *et al*., 2009; Hiwasa-Tanase *et al*., 2011; Schaart *et al*., 2011; Li *et al*., 2012; Limkul *et al*., 2015; Diamos and Mason, 2018; Pérez-González and Caro, 2018; Rosenthal, 2018; de Felippes *et al*., 2020; Wang *et al*., 2020; de Felippes *et al*., 2022).

The distinct impact of different terminators on transgene expression is not entirely clear. In most cases, the terminator-dependent increase in gene expression was correlated with higher mRNA steady-state levels, indicating that the effect of terminators on gene expression might occur at the transcriptional level and/or affect mRNA stability (Dean *et al*., 1989; Ingelbrecht *et al*., 1989; Chen *et al*., 1998; Richter *et al*., 2000; Ali and Taylor, 2001b; Nagaya *et al*., 2009; Yang *et al*., 2009; Hirai *et al*., 2011; Hiwasa-Tanase *et al*., 2011; Schaart *et al*., 2011; Li *et al*., 2012; Kurokawa *et al*., 2013; Rosenthal, 2018; de Felippes *et al*., 2020, 2022; Wang *et al*., 2020). Supporting this scenario, terminators leading to strong transgene expression showed lower levels of read-through transcription than those resulting in weaker expression (Hiwasa-Tanase *et al*., 2011; Rosenthal, 2018; de Felippes *et al*., 2020, 2022). One of the consequences of read-through transcription is the generation of aberrant transcripts, such as those lacking a poly (A) tail. Unadenylated mRNAs tend to accumulate in the nucleus, therefore not being translated, and are more susceptible to degradation by the action of exoribonucleases (de Felippes and Waterhouse, 2022).

Another consequence of read-through transcription is gene silencing. Transcripts lacking a poly (A) tail were shown to be targeted by the RNA-DEPENDENT RNA POLYMERASE 6 (RDR6) (Luo and Chen, 2007; Baeg *et al*., 2017; Sakurai *et al*., 2021), which converts single-stranded RNA into double-stranded RNA (dsRNA), the precursor molecule of small RNAs (sRNAs) (Willmann *et al*., 2011). Interestingly, transgenes are particularly susceptible to silencing mediated by sRNAs, which can reduce or completely extinguish transgene expression (Rajeevkumar *et al*., 2015; de Felippes and Waterhouse, 2020). Several reasons have been proposed to explain the high susceptibility of transgenes to silencing, including the presence or nature of different genetic elements (de Felippes and Waterhouse, 2020). In line with the link between transcription termination and silencing, the choice of terminator is a key factor affecting the production of sRNAs from transgenic transcripts in transient expression systems (de Felippes *et al*., 2020, 2022).

Introns are another example of a genetic element that can significantly impact the levels of transgene expression. In addition to enabling translation of multiple proteins from a single gene by exon shuffling, introns have been used to boost transgene expression in a phenomenon referred to as intron-mediated enhancement (IME). Although the mechanisms leading to IME are not fully elucidated, it is believed that introns may affect transgene expression by enhancing transcription, translation and/or mRNA stability (Laxa, 2017; Shaul, 2017). Introns were also shown to protect transgenes from becoming silenced. Adding introns to a transgene sequence effectively reduces RDR6-dependent gene silencing in *Arabidopsis thaliana* (Christie *et al*., 2011). In *Nicotiana benthamiana*, transgenes containing introns were also shown to be less prone to silencing; however, the specific contribution of introns to this effect was not analysed (Dadami *et al*., 2013, 2014).

We previously reported a comprehensive analysis of the contributions of different genetic elements to the expression levels and protection of transgenes against silencing (de Felippes *et al*., 2020). Using a transient expression system based on agroinfiltration of *N. benthamiana* leaves, we demonstrated that terminators were the primary determinants of transgene performance, both by suppressing sRNA production and enhancing transgene expression. In contrast to previous studies, introns did not reduce the accumulation of transgene-derived sRNAs, and their effect on GFP (GREEN FLUORESCENT PROTEIN) expression was dependent on the terminator used. Here, we extend these findings by examining the contributions of different terminators and intron presence to the stable expression of transgenes in plants. Consistent with the transient expression system, terminator choice had a major impact on transgene expression levels. This effect was associated with the efficiency of transcription termination; however, unlike in the transient system, it was not linked to sRNA-mediated silencing. Our results further reveal a strong relationship between transgene expression level and expression stability and confirm an important role for introns in protecting transgenes from silencing in stable transgenic lines. Finally, genome-wide analyses revealed that intron-containing genes are enriched among highly expressed endogenous genes and among genes showing strong transcriptional responses to stress, suggesting that protection from silencing may have contributed to the emergence and evolutionary retention of intron-containing genes.

## Results

### The effect of terminators on stable transgene expression is linked to transcriptional termination and is independent of the activity of sRNAs

To investigate how terminators and introns contribute to transgene expression levels and protection against silencing in stably transformed plants, we generated *A. thaliana* transgenic lines expressing some of the constructs used in our previous study, which focused on a transient expression system (de Felippes *et al*., 2020). These constructs consist of the RuBisCO small subunit (RBCS1A, AT1G67090) promoter (pRBCS) from *A. thaliana,* followed by the *GFP*, or a modified version thereof containing the two introns of RBCS1A (GiFiP) as a reporter gene, in combination with the terminator of one of the following genes: *NOPALINE SYNTHASE* (*NOS*) from *Agrobacterium tumefaciens*; the *RBCS1A*; or the *HEAT SHOCK PROTEIN18.2* (*HSP18.2*, AT5G59720), both from *A. thaliana* (tNOS, tRBCSA and tHSP, respectively) (Figure 1a). Analysis of 3-week-old T1 independent lines reiterates the importance of terminator choice in the final levels of transgene expression (Figure 1b-c). GFP fluorescence was higher when the transgene was expressed under the regulatory control of tHSP, while no statistically significant differences were observed with the NOS or RBCS terminators. GFP expression with tRBCS, however, was less variable than that with tNOS. We also observed an effect of introns on the final levels of transgene expression, but this was only evident when GiFiP was combined with the tHSP (Figure 1c).

**Figure 1.**
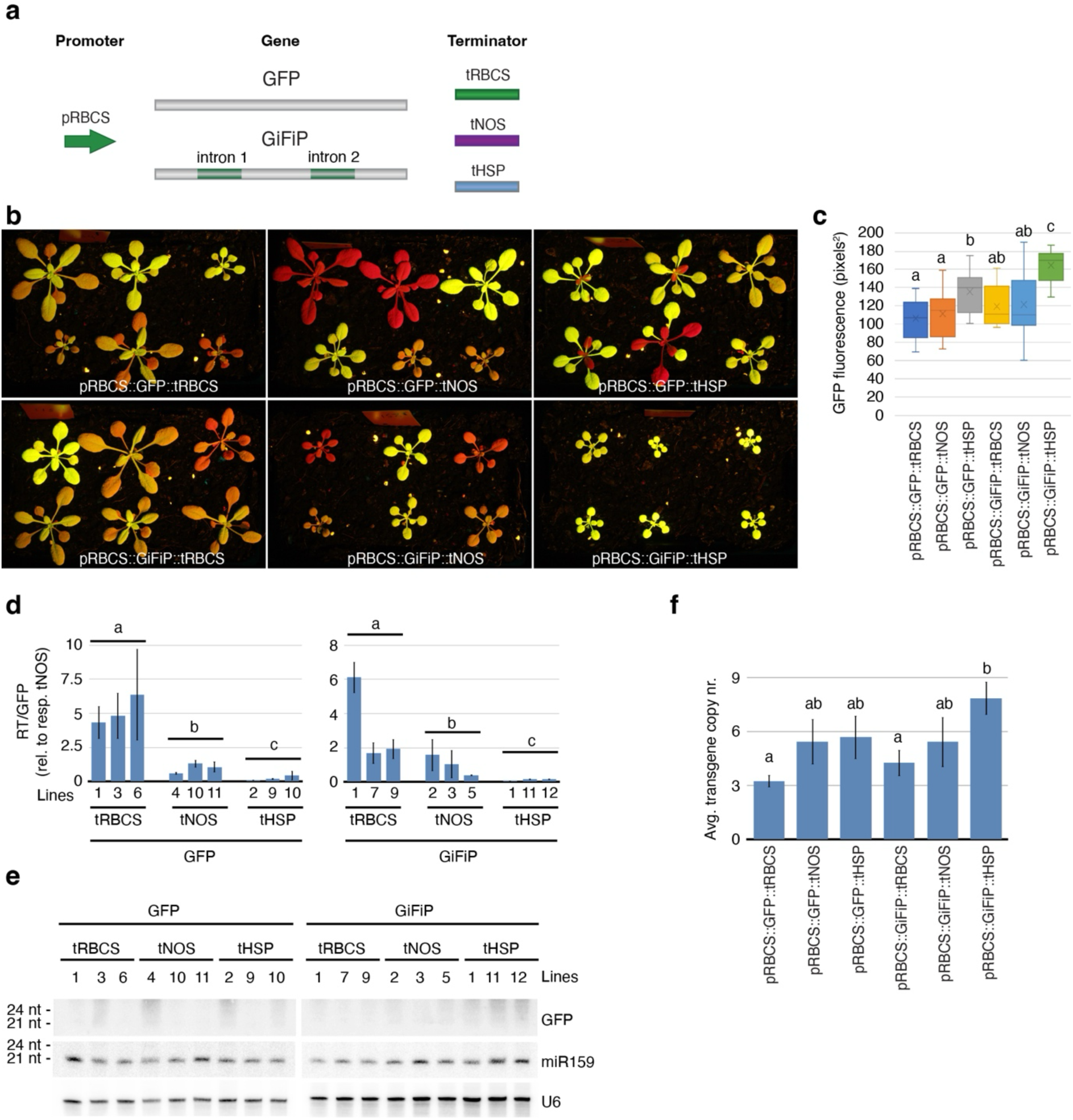
Effect of introns and different terminators on transgene expression in stable transformed plants. (a) Scheme of the different genetic elements used to generate the reporter constructs. (b) T1 *A. thaliana* plants illustrating the GFP phenotype obtained with the different constructs tested. (c) GFP fluorescence of 12 independent T1 plants. (d) RT-qPCR showing the ratio of read-through transcripts to the total GFP molecules. For each construct, three independent lines were tested. (e) The presence of sRNAs originating from the GFP transgene was tested by sRNA northern blots. miR159 and U6 were used as sRNA and loading control, respectively. RNA was extracted from a pool of T2 seedlings (11 days old) of three independent lines. (f) Average transgene copy number of 12 independent lines as detected by ddPCR. For panels c, d and f, standard errors are given, and the significance of differences was calculated using the Kruskal-Wallis test followed by the Conover procedure adjusted by the Benjamini-Hochberg FDR method (p ≤ 0.05).

We previously showed that the effects of different terminators on the GFP transient expression were linked to their transcription termination efficiency and to the accumulation of sRNAs from the transgene (de Felippes *et al*., 2020). We have therefore analysed these two aspects using quantitative reverse transcriptase-polymerase chain reaction (RT-qPCR) and Northern blot to detect read-through transcription and sRNAs, respectively. We observed a clear correlation between low levels of read-through transcription and high GFP expression, indicating the importance of efficient transcription termination for transgene expression (Figure 1d). In contrast, we did not detect transgene-derived sRNAs in any of the lines tested (Figure 1e), suggesting that silencing is not involved in the differences in GFP expression observed with the different terminators.

An important aspect that may influence the final levels of GFP expression is the number of transgene copies inserted into the genome. For half of the constructs, we can detect a correlation (R^2^ ≥ 0.7) between transgene copy number and GFP fluorescence within lines expressing the same construct (Figure S1). Variation in copy number could also explain some of the differences observed between constructs, but it is unlikely to account for all the detected variation. For instance, plants expressing pRBCS::GFP::tNOS and pRBCS::GFP:tHSP show distinct GFP fluorescence yet carry the same average number of insertions (Figure 1f).

### Introns inhibit silencing associated with high levels of transgene expression

Despite high levels of transgene expression achieved with tHSP, several 3-week-old plants carrying the pRBCS::GFP::tHSP construct exhibited signs of silencing that began later in their development and affected younger leaves (Figures 1b and 2a). Silencing was also present in shoots and siliques, but not the seeds, indicating that silencing was limited to post-transcriptional gene silencing, which is supported by the accumulation of mainly 21 nt long sRNAs (Figure 2a-c). In contrast, silencing was absent or significantly delayed in plants expressing pRBCS::GiFiP::tHSP, corroborating previous observations that introns can confer protection against silencing to transgenes that are stably expressed (Christie *et al*., 2011). Interestingly, silencing in pRBCS::GFP::tHSP lines was observed only in plants with high GFP expression, suggesting that a threshold must be exceeded for silencing to be triggered (Figure 2d). For pRBCS::GFP::tHSP, this threshold was approximately 140 pixels^2^ for plants showing early or late silencing (when silencing is visible before or after 4 weeks from germination, respectively). For plants transformed with pRBCS::GiFiP::tHSP, we only observed late silencing, and only for plants displaying GFP fluorescence higher than 174 pixels^2^, indicating an effect of introns in raising this putative threshold. The only line exhibiting late silencing and detectable sRNA accumulation outside the tHSP background was line #3 carrying the pRBCS::GFP::tNOS construct. Notably, this was also the only line among the pRBCS::GFP::tRBCS and pRBCS::GFP::tNOS transformants with GFP fluorescence exceeding the proposed threshold of 140 pixels², further supporting the hypothesis that silencing is triggered once transgene expression surpasses a critical level (Figure S2).

**Figure 2.**
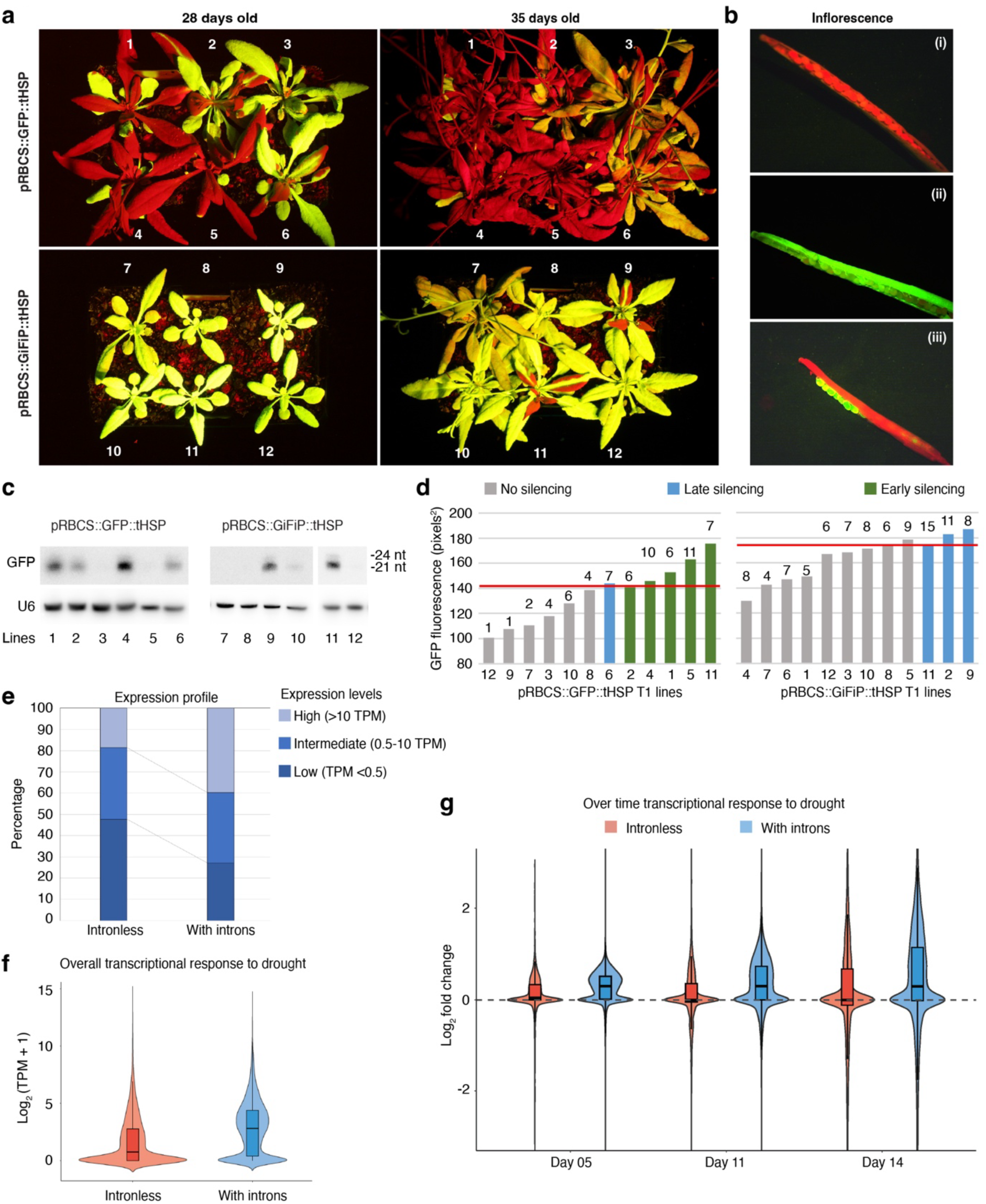
Silencing inhibition and gene expression boost mediated by introns. (a) Representation of silencing affecting some of the plants with high transgene expression and the inhibitory effect of introns on this phenomenon. Plants depicted here are the same shown in Figure 1. (b) Detail of the siliques of wild-type (i), GFP-expressing (ii) and GFP-silenced plants (iii) showing that GFP expression in seeds is not affected by silencing. (c) sRNA accumulation in the plants shown in panel (a) as detected by sRNA northern blot using floral tissue. U6 is used as a loading control. (d) Correlation between GFP expression level and the onset of silencing. The red line represents the GFP expression threshold for silencing to be triggered. Early or late silencing applies to silencing visible before or after 4 weeks from germination, respectively. Transgene copy number for each line is provided at the top of their respective column. (e) Expression profile of intronless and intron-containing genes. Overall (f) and over time (g) transcriptional response of intronless and intron-containing genes to drought stress.

### Introns are associated with elevated expression and enhanced stress responsiveness of endogenous genes

The idea that a threshold needs to be reached for triggering transgene-dependent sRNA production, and the role introns seem to have in this process, led us to the attractive hypothesis that one of the forces resulting in the emergence of introns was the possibility for genes to achieve higher levels of expression without becoming targets for sRNAs. In this case, one would expect to find a prevalence of intron-containing genes among the highly expressed genes in a plant genome. To test this hypothesis, we compared the expression profiles of intronless and intron-containing genes in *A. thaliana*. Intron-containing genes exhibited significantly higher expression than intronless genes, as supported by the Mann-Whitney U test (p-value < 0.001) (Figure 2e). When gene expression was classified into low (TPM <0.5), intermediate (0.5-10 TPM), and high (>10 TPM) expression categories based on expression quantiles, a distinct distribution pattern was observed. Intronless genes were disproportionately overrepresented in the low-expression category (47%), whereas only 18% were found in the high-expression category. In contrast, 40% and 27% of intron-containing genes were enriched in the high- and low-expression categories, respectively. Building on these differential expression profiles, we subsequently evaluated which genes exhibited the most pronounced transcriptional response to stress. Using a time-course of water deficit data for *A. thaliana* (Day 00, Day 05, Day 11, and Day 14) (Rosa *et al*., 2019), we analysed changes in gene expression to determine whether intron presence influences the magnitude of stress-responsive transcriptional activation or suppression. Across all time points, comparison of log_2_ fold change values showed that intron-containing genes tended to maintain stable expression or become induced under stress, whereas intronless genes were more often unable to show induction (Figure 2f-g). These genome-wide trends corroborate the idea that introns, as in transgenic systems, confer an advantage for highly expressed endogenous genes, presumably by raising the expression threshold that triggers silencing. Consistent with this hypothesis, genome-wide analyses conducted elsewhere have shown that intronless genes are disproportionately represented among the sources of endogenous sRNAs (Christie *et al*., 2011), supporting the idea that one of the evolutionary drivers behind the evolution of introns was the prevention of silencing in highly expressed genes.

## Discussion

In this work, we complement a previous study from our group, in which we analysed the contribution of terminators and introns to the stability and final expression levels of transgenes. As in transient expression systems, terminators were key elements affecting transgene expression in stable transformants. Each of the terminators tested had a distinct effect on the final GFP expression levels, with a pattern similar to that observed in the agroinfiltration assay (de Felippes *et al*., 2020). Consistently, transgene expression was once again linked to the efficiency of each terminator in promoting transcription termination. In contrast, although sRNA-mediated silencing was detected in some lines at later developmental stages, it did not contribute to the variation in transgene expression observed with the different constructs in younger, stably transformed plants. This observation indicates that differences in the final levels of stable transgene expression were mainly due to unproductive transcription and post-transcriptional events mediated by the terminator. In this scenario, poor transcription termination results in read-through transcription, which is the source of aberrant transcripts, such as unadenylated ones (Luo and Chen, 2007). The poly (A) tail is an important feature for the stability, nuclear export and translation of mRNAs, and consequently, its absence has a negative consequence for protein expression. Moreover, aberrant transcripts are mostly degraded by the RNA decay pathway before they can be translated into proteins, affecting the final expression levels of a gene (Mandel *et al*., 2007; Hung and Slotkin, 2021; de Felippes and Waterhouse, 2022).

Despite not being involved with the differences in transgene expression levels observed among distinct constructs, silencing was detected in several plants later in their development. Although most of these plants were dependent on the tHSP function, this late silencing was strongly associated with intense GFP fluorescence, suggesting that it was due to high transgene expression levels rather than poor terminator efficiency. In accordance, high expression is one of the factors associated with transgene silencing (de Felippes and Waterhouse, 2020). Moreover, our data point to the existence of a threshold that needs to be exceeded to induce sRNA production from transgenes (Figure 2d and S2). The existence of an expression threshold for silencing is supported elsewhere, for example, by the work of Schubert and colleagues (Schubert *et al*., 2004) with transgenic petunias. Analysis of a transgenic population showed a correlation between high transgene expression and transgene copy number, and that silencing is more likely to happen when a certain number of insertions are present. This expression threshold might reflect the dominance of the RNA decay machinery over the sRNA pathway in processing aberrant transcripts. Accordingly, suppression of the RNA decay mechanism results in increased levels of sRNAs originating from endo- and transgenes (Gazzani *et al*., 2004; Gregory *et al*., 2008; Moreno *et al*., 2013; Branscheid *et al*., 2015; Yu *et al*., 2015; Zhang *et al*., 2015). In this scenario, aberrant transcripts would be primarily degraded by the RNA decay machinery. Higher levels of expression could result in aberrant transcript levels that saturate this degradation pathway, thereby giving RDR6 access to such transcripts and initiating sRNA production.

As for transient expression, the presence of introns alone was insufficient to boost transgene expression in stable lines. However, when combined with the tHSP, introns significantly contributed to the overall transgene expression levels. Given the lack of sRNAs in young plants, this contribution might have occurred through a mechanism related to the IME phenomenon (Laxa, 2017; Shaul, 2017). We also observed in lines prone to silencing an inhibitory effect of introns on the generation of transgene-derived sRNAs. This protective role of introns against silencing was previously shown for stably transformed plants, but it was not detected in our previous study when the same constructs used here were transiently expressed (Christie *et al*., 2011; de Felippes *et al*., 2020). Splicing might facilitate the routing of aberrant transcripts to the RNA decay pathway, outcompeting RDR6 for access to these molecules (Christie *et al*., 2011). Given that introns can have a positive effect on transcription termination (Dwyer *et al*., 2021), it is also possible that the intron-dependent inhibition of silencing occurs through the reduction of read-through events and aberrant transcript generation from transgenes.

Regardless of the mechanism, our data suggest that the intron protection against silencing allows for higher gene expression by increasing the threshold associated with the onset of silencing. This feature might have been one of the evolutionary forces behind the rise of introns in plants, allowing robust gene expression without triggering silencing. This hypothesis is supported by genome-wide analysis showing the preponderance of intronless genes as the source of endogenous sRNA populations (Christie *et al*., 2011).

According to the Gene Harmony model, optimal transgene expression is achieved when terminators counterbalance promoter-driven transcription, thereby minimising read-through transcription and the production of sRNAs (de Felippes *et al*., 2020, 2022; de Felippes and Waterhouse, 2022). Based on the findings presented here, we propose an updated version of this model that better accounts for the differences observed between transient and stable transgenic systems (Figure 3). In transient expression systems, transgene expression originates from both genome-integrated and extrachromosomal double-stranded T-DNAs (dsT-DNAs) (Lacroix and Citovsky, 2013; Thomson *et al*., 2024). Given the high abundance of dsT-DNAs within plant cells (Janssen and Gardner, 1990; Buck *et al*., 2000), transgene transcripts are produced at exceptionally high levels, often exceeding the threshold at which silencing is triggered due to saturation of RNA decay pathways. Under these conditions, differences in terminator performance become readily apparent, with strong terminators enhancing transgene expression by limiting read-through events and decreasing sRNA accumulation, consequently reducing unproductive transcription and inhibiting silencing (Figure 3a-b). In contrast, transgene expression in stably transformed plants derives exclusively from T-DNA copies integrated into the plant genome, substantially reducing the number of transcripts produced per cell. As a result, transcript levels are less likely to exceed the threshold required to trigger silencing. In this context, aberrant transcripts are predominantly removed by the RNA decay machinery, and the effect of terminators on transgene expression is largely restricted to their influence on unproductive transcription (Figure 3c-d). However, in some cases, such as plants carrying multiple T-DNA insertions, transcript abundance may surpass the threshold that induces silencing. Under these circumstances, the situation resembles that of transient expression systems, where inefficient terminators generate large amounts of aberrant transcripts that overwhelm RNA quality-control mechanisms, leading to elevated sRNA production, strong silencing, and reduced transgene expression (Figure 3e). By contrast, strong terminators minimise read-through transcription and the generation of aberrant RNAs, delaying the onset of silencing. Nevertheless, silencing may still arise in specific tissues or developmental stages, resulting in strong yet unstable transgene expression over time (Figure 3f). Introns serve as an additional layer of protection against silencing, inhibiting sRNA production from highly expressed transgenes through a mechanism that is still not entirely understood (Figure 3g).

**Figure 3.**
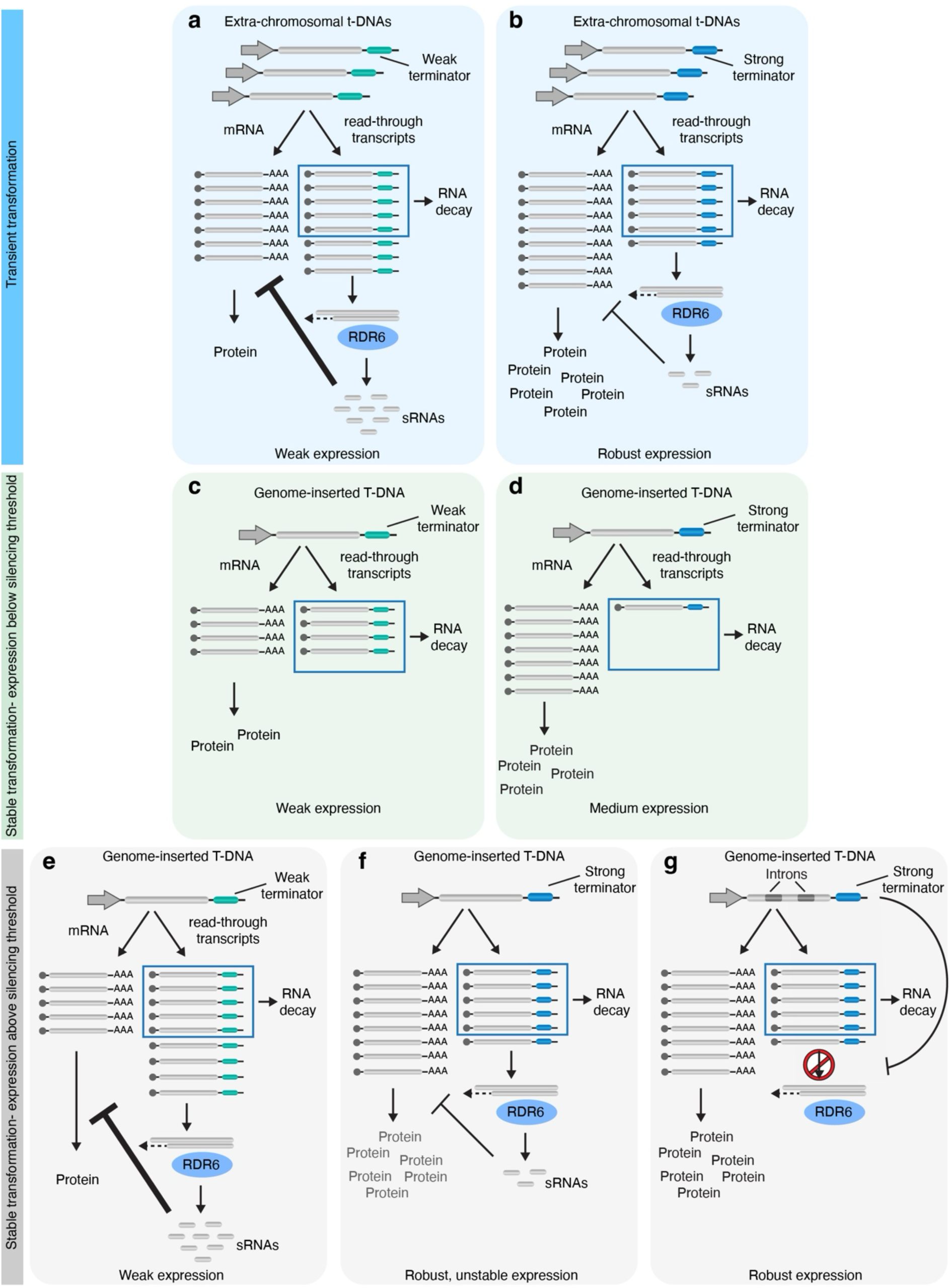
Gene Harmony model for the role of terminators in the expression of transgenes in plants. In transient expression, the numerous T-DNAs present as extrachromosomal molecules lead to an elevated number of transcripts. (A) When the expression involves the deployment of a weak terminator, a significant number of aberrant transcripts is produced, surpassing the capacity of the RNA decay machinery and triggering abundant sRNA production that severely affects final transgene expression. (B) Efficient terminators result in fewer aberrant transcripts and consequently, in an increase of regular mRNAs that are translated and less silencing due to a lower level of sRNAs. (C) In stable transformants, despite the relatively high levels of read-through transcription resulting from the poor performance of a weak terminator, the reduced number of T-DNAs being transcribed prevents saturation of the RNA decay pathway and stops RDR6 from accessing aberrant transcripts. Nonetheless, transgene expression is compromised by a low number of regular mRNAs due to relatively high read-through transcription. (D) Efficient terminators contribute to final expression levels by improving the ratio of regular mRNAs to aberrant transcripts. (E) In cases where transcript levels are elevated (e.g. due to the presence of multiple T-DNA copies), the number of aberrant transcripts might exceed the capacity of the RNA decay machinery, resulting in sRNA production, especially when weaker terminators are used. (F) For strong terminators, sRNA accumulation would be reduced or restricted to certain tissues and/or developmental stages, resulting in stronger but unstable expression. (G) The presence of introns can bring stability to highly expressed transgenes by inhibiting RDR6 activity through a still unknown mechanism.

The findings presented here further emphasise the importance of genetic elements beyond promoters in determining transgene expression. They demonstrate that optimal expression emerges from the combinatorial action of multiple regulatory components and provide new insights into the mechanisms through which terminators and introns influence transgene performance.

## Material and Methods

### Plant material and transformation

*A. thaliana* plants (Col-0) were grown at 23°C under long days condition (16 h light/ 8 h dark). Plants were transformed with the floral dip method (Clough and Bent, 1998), and transgenic plants were selected on ½ MS plates supplemented with 12 mg/L of Glufosinate-ammonium PESTANAL® (Sigma-Aldrich) and 100 mg/L of Carbenicillin.

### GFP images and quantification

GFP fluorescence was visualised using blue light from a Dark Reader Hand Lamp (HL32T; Clare Chemical Research, Dolores, CO, USA) in combination with an orange filter. Images were captured using an EOS 550D camera (Canon, Tokyo, Japan) fitted with an Orange G HMC filter (HOYA Filters, Tokyo, Japan), using manual settings adjusted to each experiment as required. GFP fluorescence was quantified using FIJI software (Schindelin *et al*., 2012). Images of 19-day-old plants were first converted to 16-bit format. A circular region of interest (width = 36, height = 36, area = 1020) was then defined, and the mean grey value (i.e., the intensity of a pixel in a grayscale image) was obtained using the Measure tool. For each plant, the reported grey value represents the average of three independent measurements of the 3^rd^/4^th^ true leaf.

### RNA extraction, sRNA Northern blots and RT-qPCR

Total RNA was extracted with TRIzol reagent (Thermo Fisher Scientific, Waltham, MA, USA). Small RNA species were examined by sRNA Northern blot analysis. Briefly, 4–6 µg of total RNA was separated on a 17% denaturing polyacrylamide gel containing 7 M urea, transferred onto a positively charged membrane, and immobilised by UV crosslinking before hybridisation with specific probes (Table S1). Both random-primed and oligonucleotide DNA probes were labelled with α-³²P-dCTP using either the Prime-a-Gene Labelling System (Promega, Madison, USA) or terminal deoxyribonucleotidyl transferase (Fermentas, Thermo Fisher Scientific), respectively.

RT-qPCR analysis to detect transcriptional read-through was performed using 80 ng of total RNA extracted from T2 plants. For each line, three individual 11-days-old seedlings were used to calculate the average read-through ratio. RNA samples were treated with RQ1 DNase (Promega, Madison, WI, USA) and subsequently reverse-transcribed into cDNA using SuperScript III (Invitrogen, Carlsbad, CA, USA) in the presence of oligo(dT) and a specific primer annealing downstream of the terminator (RT primer). qPCR reactions were prepared with GoTaq qPCR Master Mix (Promega), 10 ng of cDNA, and 10 pmol of either gene-specific primers or read-through primer pairs, in a final volume of 10 μl. Amplification was carried out on a CFX384 Real-Time PCR System (Bio-Rad, Hercules, CA, USA). Cycling conditions consisted of an initial denaturation at 95 °C for 10 min, followed by 39 cycles of 95 °C for 15 s and 60 °C for 30 s. Melting curve analysis was performed by increasing the temperature from 65 °C to 95 °C in 0.5 °C increments, with a 5 s hold at each step. *ACTIN* served as the reference housekeeping gene. Primer sequences are listed in Table S1.

### Transgene copy number estimation by ddPCR

To determine the number of T-DNA copies integrated in different transgenic lines, a duplex probe-based ddPCR assay was performed using the QX200 Droplet Digital PCR System (BioRad, Hercules, CA, USA). Genomic DNA was extracted from leaves of T1 plants using the CTAB method and digested with HindIII (New England BioLabs, Ipswich, MA, USA) for 1 h at 37°C. PCR reaction was prepared in 20 µl total volume, using 250 nM of probe, 450 nM for each primer, 25 ng of digested genomic DNA and ddPCR Supermix for Probes (BioRad). HMGB1 (AT3g51880) was used as the single-copy reference gene (Ruiz-Salas *et al*.), and copy number determination was based on quantification of the selectable marker gene phosphinothricin acetyltransferase. Droplet generation, PCR conditions and droplet reading were performed according to the manufacturer’s instructions. Primers and probes sequences are given in Table S1.

### RNA-seq data analysis

RNA-seq data set from *A. thaliana* water deficit time-course experiment (Rosa *et al*., 2019) were retrieved from the NCBI (SRA) under the following accessions; Day 00 (SRR7624718, SRR7624719, SRR7624720), Day 05 (SRR7624715, SRR7624716, SRR7624717, SRR7624700, SRR7624701, SRR7624695), Day 11 (SRR7624710, SRR7624714, SRR7624723, SRR7624694, SRR7624696, SRR7624697), and Day 14 (SRR7624698, SRR7624699, SRR7624711, SRR7624706, SRR7624707, SRR7624711). Bioinformatics data analysis was performed on the Galaxy Australia platform (Community *et al*., 2024). Raw reads were processed with Trimmomatic (Bolger *et al*., 2014) to remove adapters and low-quality bases, and transcript quantification was performed with Kallisto (Bray *et al*., 2016). Differential gene expression analysis was conducted using DESeq2 (Love *et al*., 2014), and genes were subsequently classified into expression categories based on 33% quantile calculations to define low, intermediate, and high expression tiers. Statistical significance between groups was assessed using the Mann-Whitney U test, and all data visualisations, including expression profiles and comparative plots, were generated using the ggplot2 (Wickham, 2016) package in R Studio (R. Posit Software, PBC, Boston, MA).

## Supplementary data

Table S1. List of primers and probes.

Figure S1. Effect of transgene copy number on GFP expression.

Figure S2. Late silencing in plants carrying the tRBCS and tNOS.

## Acknowledgements and funding

This work was funded by the Australian Research Council (ARC) grant number CE200100015 and FL160100155.

## Author contributions

FFF designed the research; BR, FFF and KS performed research; BR, FFF and PMW analysed the data and BR and FFF wrote the manuscript.

