## Supplementary Figures and Tables for "The combinatorial effect of terminators and introns on the levels and stability of stable transgene expression in plants"

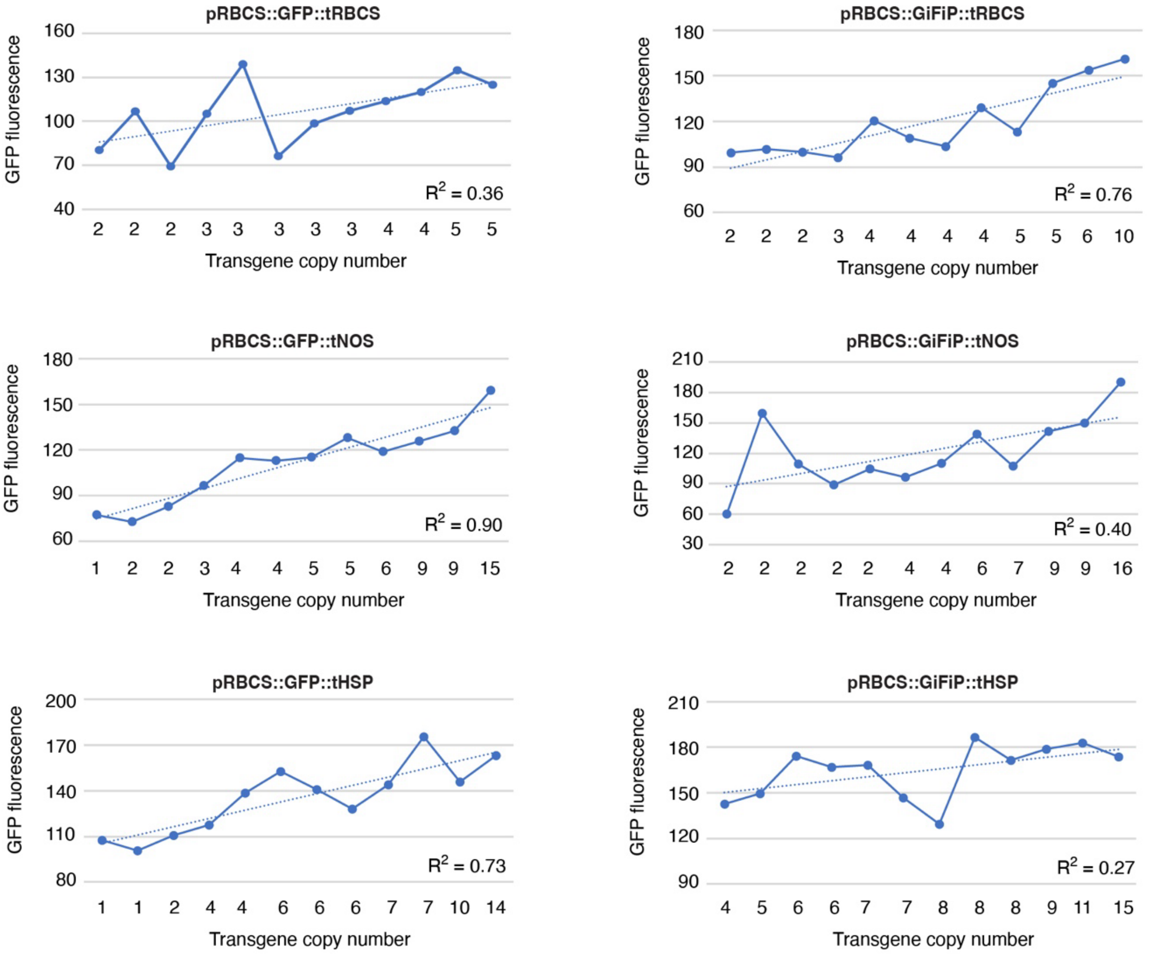


**Figure S1. Effect of transgene copy number on GFP expression.** The correlation between GFP fluorescence (pixels$^2$) and transgene copy number for each independent transgenic line is shown. The coefficient of determination ($R^2$) for each construct is given.

Figure S2

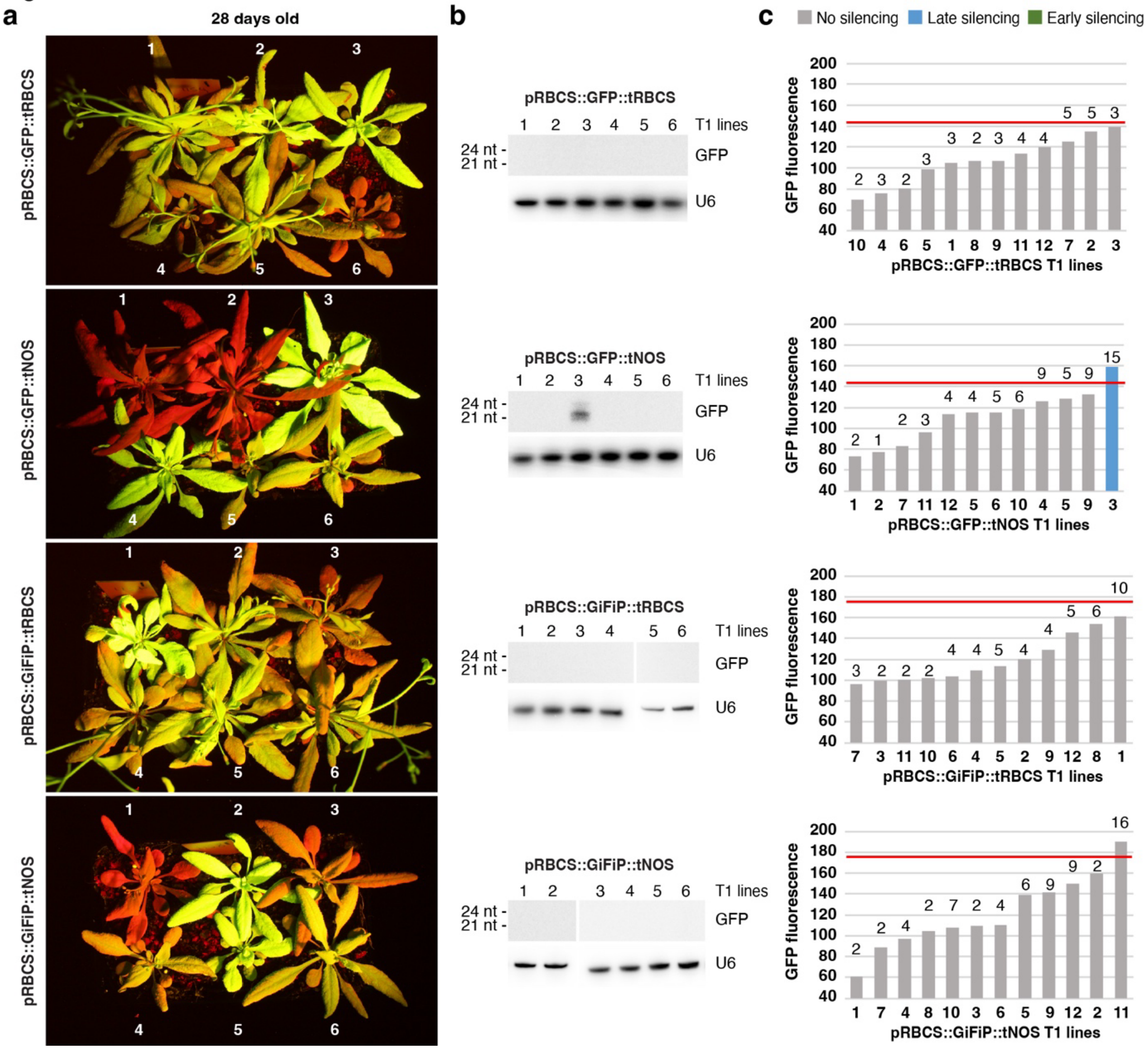


**Figure S2. Late silencing in plants carrying the tRBCS and tNOS.** (a) Plants representing the onset of late silencing in plants carrying the tRBCS and tNOS. These are the same plants shown in Figure 1. (b) sRNA northern blot to detect sRNAs originating from the GFP sequence. (c) Correlation between GFP fluorescence (pixels$^2$) and silencing. Red lines represent the expression threshold triggering silencing as defined by lines carrying the tHSP (Figure 2d). The number at the top of each column represents the number of transgene insertions for each line. Early or late silencing is defined by the presence of silencing occurring before or after 4 weeks from germination, respectively.

**Table S1.** List of primers and probes used in this work

| Primer name | sequence | target | Observation |
|---|---|---|---|
| ***RT-qPCR*** | | | |
| FQ-102 | TCCCGACTGATGTCAGAGC | HPL1 promoter | region downstream of terminator. Specific primer for cDNA synthesis |
| FQ-103 | CGTGGATACTTGGCAGTGG | HPL1 promoter | Read-through primer forward |
| FQ-104 | TGTTGTAGCGTTACTATGAAGACC | HPL1 promoter | Read-through primer reverse |
| FQ-881 | AGCAAAGACCCCAACGAGAA | GFP | |
| FQ-004 | CGTGGATACTTGGCAGTGG | GFP | |
| FQ-105 | GCACCCTGTTCTTCTTACCG | ACTIN | Housekeeping gene |
| FQ-106 | AACCCTCGTAGATTGGCACA | ACTIN | Housekeeping gene |
| ***sRNA blot probes*** | | | |
| FQ-003 | AGATCCGCCACAACATCGAG | GFP | |
| FQ-004 | TTGTACAGCTCGTCCATGCC | GFP | |
| FQ-107 | AGGGGCCATGCTAATCTTCTC | U6 | |
| miR159 | TAGAGCTCCCTTCAATCCAAA | miR159 | |
| ***ddPCR*** | | | |
| FQ-162 | AGCCACAAACACCACAAGA | phosphinothricin acetyltransferase | |
| FQ-163 | AGCAATACCAGCCACAACA | phosphinothricin acetyltransferase | |
| FQ-146 | TACCCTTGGTTGGTTGCTGAGGTT | phosphinothricin acetyltransferase | Probe for ddPCR. It has a 5' HEX and 3' IABkFQ |
| FQ-156 | GTGCTTGCTTTCTGAAGATGTATG | HMGB1 | |
| FQ-157 | CGTCATCTTCTTCCTCTTCTTCTT | HMGB1 | |
| FQ-149 | ATCACCTGCCGCTTCCTTCT | HMGB1 | Probe for ddPCR. It has a 5'-FAM and 3' IABkFQ |
